# Dual oxic-anoxic co-culture enables direct study of host-anaerobe interactions at the airway epithelial interface

**DOI:** 10.1101/2021.03.05.433759

**Authors:** Patrick J. Moore, Talia D. Wiggen, Leslie A. Kent, Sabrina J. Arif, Sarah K. Lucas, Joshua R. Fletcher, Alex R. Villareal, Adam Gilbertsen, Scott M. O’Grady, Ryan C. Hunter

**Author notes:** To whom correspondence should be addressed: Ryan C. Hunter, Department of Microbiology & Immunology, Microbiology Research Facility, 3-115 University of Minnesota, 689 23^rd^ Avenue SE, Minneapolis, MN 55455, Twitter: @hunter_lab.

## Abstract

Strict and facultative anaerobic bacteria are widely associated with both acute and chronic airway disease. However, their potential role(s) in disease pathophysiology remain poorly understood due to inherent limitations of existing models and conflicting oxygen demands between anaerobes and host cells. To overcome these limitations, we optimized a dual oxic-anoxic culture (DOAC) platform that maintains an oxygen-limited microenvironment at the epithelial interface while host cells are oxygenated basolaterally. This approach enables host-bacterial co-culture for ∼24h, and here we demonstrate its utility via two applications. First, we show that anaerobe challenge results in epithelial mucus degradation, inflammatory marker gene expression, and enhanced pathogen colonization. Second, we combine DOAC with single cell RNA sequencing (scRNAseq) to reveal a cell type-specific transcriptional response of the airway epithelium to anaerobe infection. Together, these data illustrate the versatility of DOAC while revealing new insights into anaerobe-host interactions and their mechanistic contributions to airway disease pathophysiology.

## INTRODUCTION

Decades of clinical laboratory culture have focused on a limited set of pathogens associated with acute and chronic airway disease (e.g., *Pseudomonas aeruginosa, Staphylococcus aureus, Mycobacterium tuberculosis*). More recently, culture-independent sequencing studies of airway microbiota have identified more complex bacterial signatures, lending evidence to polymicrobial disease etiologies. Notably, oral-associated facultative and obligate anaerobes – *Fusobacterium, Prevotella, Veillonella, Streptococcus* spp. *–* which are present at low densities in the healthy respiratory tract^1-4^, are both prevalent and abundant in chronic obstructive pulmonary disease (COPD), cystic fibrosis (CF), non-CF bronchiectasis, sinusitis, tuberculosis, ventilator-associated pneumonia, and other respiratory complications^5-10^. In each instance, development of hypoxic microenvironments at the airway epithelial interface provides a niche for anaerobe proliferation, often reaching densities equal to or greater than those of canonical pathogens.

Mechanistic contributions of anaerobic bacteria to airway disease remain poorly understood though several roles have been proposed. In healthy individuals, anaerobe abundance in bronchoalveolar lavage fluid correlates with expression of proinflammatory cytokines, elevated Th17 lymphocytes, and a blunted TLR4 response, implicating a compromised first line of defense against bacterial infection^1^. Indeed, epidemiologic and *in vitro* data suggest that anaerobes may facilitate secondary colonization by canonical airway pathogens. In non-CF bronchiectasis, *Prevotella* and *Veillonella* positively correlate with Th17 cytokines and non-tuberculosis mycobacterial infection^11^. Similarly, in HIV subjects, anaerobes suppress expression of interferon gamma and IL-17A via production of short-chain fatty acids (SCFAs) and are thought to impair the host response to consequent *M. tuberculosis* colonization^9^. In CF, anaerobe-derived SCFAs increase with age and disease progression^12^, mediate excessive production of IL-8 by bronchial epithelial cells (in turn promoting neutrophil mobilization)^13^ and potentiate the growth and virulence of canonical CF pathogens *in vitro*^14-15^. Anaerobe-dominated bacterial communities in the CF airways are more commonly associated with milder disease^16^, but they also increase in abundance during acute disease flares prior to antibiotic therapy^17^, implicating anaerobes in pathogenesis.

Direct studies of anaerobe-host and anaerobe-host-pathogen interactions have been limited by a dearth of compatible laboratory methods. Animal models can poorly reflect chronic infection pathologies and/or be prohibitively expensive for high throughput analyses. As an alternative, three-dimensional (3-D) cell culture approaches have greatly expanded our knowledge of microbial-epithelial interactions^18-20^. However, incorporation of anaerobic microbiota into these models is restricted by the inherent challenge of maintaining host cell viability under oxygen-limited culture conditions. New *in vitro* approaches are needed for a more detailed understanding of anaerobic microbiota and their interactions with airway epithelium.

Here we optimized a dual oxic-anoxic culture (DOAC) platform that recapitulates an oxygen limited epithelial microenvironment thought to exist in the diseased airways^21-22^. In this platform, polarized epithelial monolayers are maintained at air-liquid interface in an anaerobic chamber while O2 and CO2 are delivered to the culture apparatus from an external source. This setup allows for maintenance of hypoxia in the apical compartment while host cells are oxygenated basolaterally. We establish the utility of this model through two *in vitro* studies of anaerobe-airway interactions. First, we use the DOAC platform to test the hypothesis that anaerobic microbiota enhance colonization of the epithelial surface by the canonical airway pathogen, *P. aeruginosa*. Second, we combine DOAC with single-cell RNA sequencing (scRNAseq) to assess the cell type-specific transcriptional response of normal human tracheal bronchial epithelial (nHTBE) cells to infection by *Fusobacterium nucleatum* – an oral commensal with pathogenic potential^23^. Through this work we not only demonstrate the power and versatility of the DOAC platform, but also offer new insight into mechanisms of pathogen colonization and potential roles of anaerobic bacteria in the onset and development of airway disease.

## RESULTS

### Optimization of a dual oxic-anoxic epithelial culture (DOAC) platform

Our primary objective was to establish and optimize a cell culture platform that facilitates study of anaerobe-host interactions (**Fig. 1A, Fig. S1**). To do so, we first cultured polarized monolayers of the adenocarcinoma cell line, Calu-3, at air-liquid interface (ALI) for 21-28 days under standard (normoxic) conditions. As shown previously^24^, polarized Calu-3s produce a distinct mucus layer on the apical surface (**Fig. S1A**), mimicking aberrant mucin accumulation associated with CF, COPD, sinusitis and other chronic airway diseases. Once polarized, cell cultures were placed in a gas-permeable multi-well plate manifold (**Fig. S1B-D, Fig. S2**), transferred to an anaerobic chamber, and mixed gas (21% O2/ 5% CO2/ 74% N2) was delivered through a chamber cable gland to the basolateral compartment of the Transwell-containing plate. This dual oxic-anoxic culture (DOAC) format achieves oxygenation of host cells while maintaining exposure of the apical surface to the oxygen-limited environment of the anaerobic chamber. Using DOAC we determined that, when compared to cell culture under normoxic and strict anoxic conditions, cytotoxicity was negligible up to 24h as determined by lactate dehydrogenase (LDH) release, with a progressive increase in cell death after 48h and 72h (**Fig. 1B)**. Viability of the immortalized RPMI 2650 nasal cell line and primary human tracheal bronchial epithelial cells (nHTBE) was similarly achieved under DOAC conditions for at least 24h (**Fig. 1C)**.

**Fig. 1.**
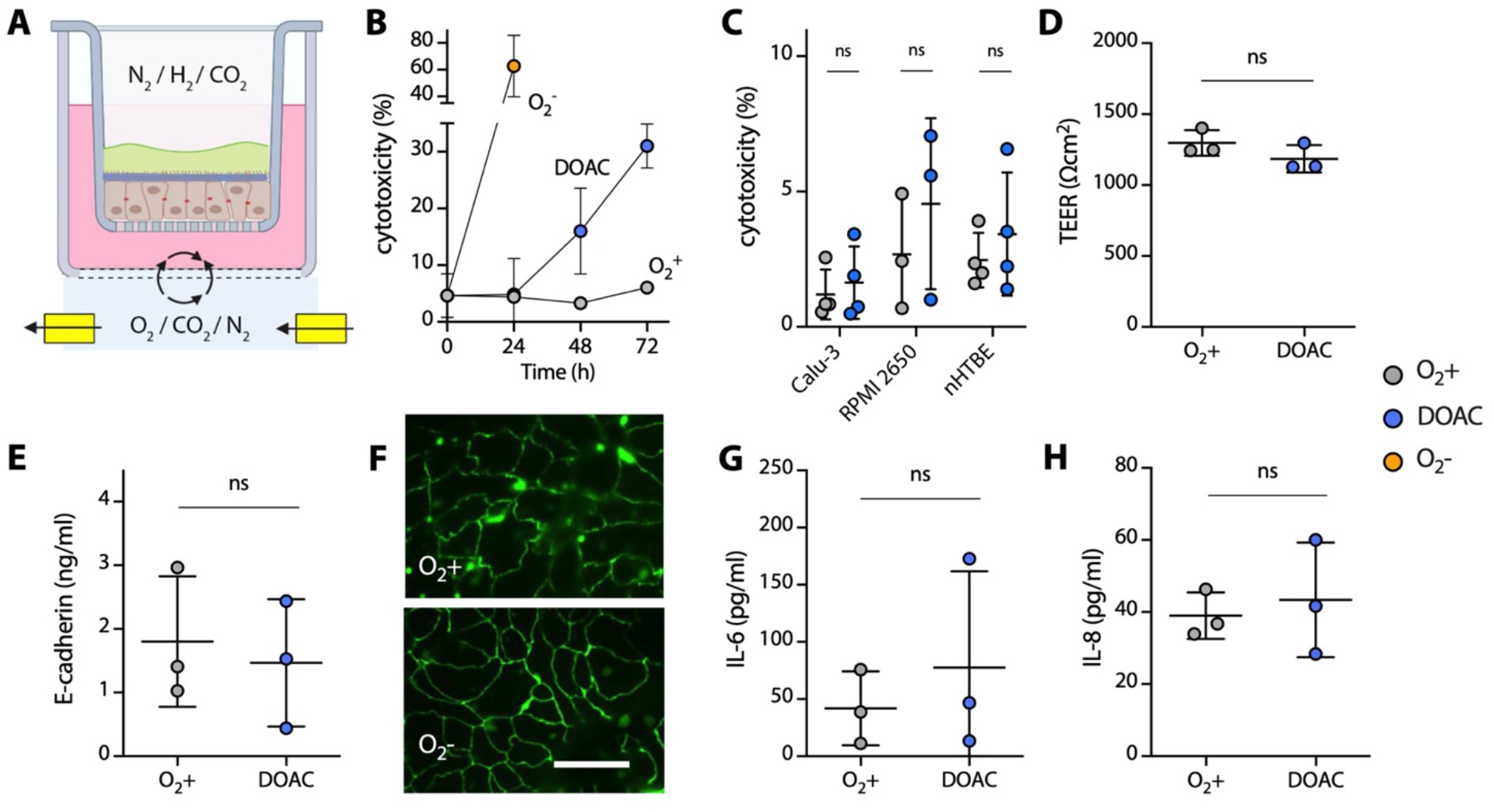
Dual oxic-anoxic culture (DOAC) of human bronchial epithelial cells. **(A)** Primary and immortalized (Calu-3, RPMI2650) cells are cultured on Transwell inserts at ALI for 21-28 days, transferred to a gas permeable multi-well plate manifold in an anaerobic chamber, and incubated under dual oxic-anoxic culture conditions where the apical compartment is oxygen-limited and mixed gas is delivered basolaterally. Figure created with BioRender.com **(B)** Cytotoxicity as measured by lactate dehydrogenase (LDH) in the DOAC culture medium over time relative to normoxic (O2+)and strict anoxic (O2-) culture conditions. LDH activity is expressed as a % relative to normoxic cultures at t=0 and a lysed control culture. **(C)** LDH release by Calu-3, RPMI 2650, and primary nHTBE cells under normoxic (grey) and DOAC (blue) culture conditions after 24h. **(D)** Transepithelial electrical resistance (TEER), **(E)** E-cadherin concentrations, **(F)** immunofluorescence of zonula occludens 1 (ZO-1) tight junction proteins, and cytokines **(G)** IL-6, and **(H)** IL-8 showed no significant differences between normoxic and DOAC culture conditions after 24h. All data shown for panels B-D, F, and G were derived from at least three independent experiments with three technical replicates each and were compared using an unpaired t-test with Welch’s correction. Images in panel F are representative of three independent experiments. Bar = 20μm.

We then determined the effects of DOAC, if any, on Calu-3 cell physiology after 24h, which was chosen as our time point for downstream experiments. Transepithelial electrical resistance (TEER) (**Fig. 1D**) and E-cadherin concentrations (**Fig. 1E**), both proxies of epithelial barrier integrity, showed slight but insignificant decreases under DOAC conditions (*P*=0.21 and 0.70, respectively). These data were corroborated by immunofluorescence microscopy which revealed confluent monolayers and well-defined staining of tight junction zonula occludens proteins that appeared as near-continuous rings at the periphery of each cell (**Fig. 1F**). Previous work has shown hypoxia-induced expression of pro-inflammatory cytokines in pulmonary fibroblasts^25^. Thus, we also used an enzyme-linked immunosorbent assay to quantify IL-6 and IL-8 production by Calu-3 cells. Both cytokines showed no significant increases under DOAC conditions relative to controls (**Fig. 1G, H**, *P*=0.54 and 0.68, respectively), suggesting that our culture conditions do not elicit an inflammatory response after 24h.

To gain a broader understanding of the physiological response of Calu-3 cells to dual oxic-anoxic culture, we used bulk RNAseq to compare global Calu-3 gene expression to culture under normoxic conditions. Transcriptome analysis revealed 148 differentially expressed transcripts (117 upregulated, 31 downregulated, l2fc β 1, adjusted *P*<0.001; out of ∼16,000 total genes) (**Fig. 2A, Data S1**). With the exception of *ANGPTL4* (encoding angiopoietin-like 4) and *SERPINA1* (alpha-1 antitrypsin) (**Fig. 2B**), which are induced in response to hypoxia and acute inflammation, respectively, few markers of cell stress were differentially expressed, including genes involved in tight junction formation, oxidative stress, and endoplasmic reticulum stress. Importantly, HIF-1α, which is constitutively expressed at low levels under normoxia but upregulated under hypoxia, was also consistent between culture conditions after 24h, suggesting that Calu-3 cells are sufficiently oxygenated under DOAC conditions (**Fig. 2B**). Among inflammatory biomarkers, only *ICAM1* (intracellular adhesion molecule 1) and *TGFβ1* (transforming growth factor beta 1) showed statistically significant differences, further demonstrating that DOAC did not yield an appreciably inflammatory microenvironment (**Fig. 2C**). Finally, since we would eventually use the DOAC model to assay bacterial colonization of the apical mucus layer, we compared mucin-related gene expression between conditions. Among detectable transcripts (*MUC1, MUC3A, MUC5AC, MUC5B*, and *MUC13*), no significant differences were observed between culture conditions (**Fig. 2D**). These data demonstrate that DOAC yields minor changes in cytotoxicity and gene expression after 24h, but we consider these changes to be negligible relative to the value of the DOAC platform for modeling anaerobe-epithelial interactions.

**Fig. 2.**
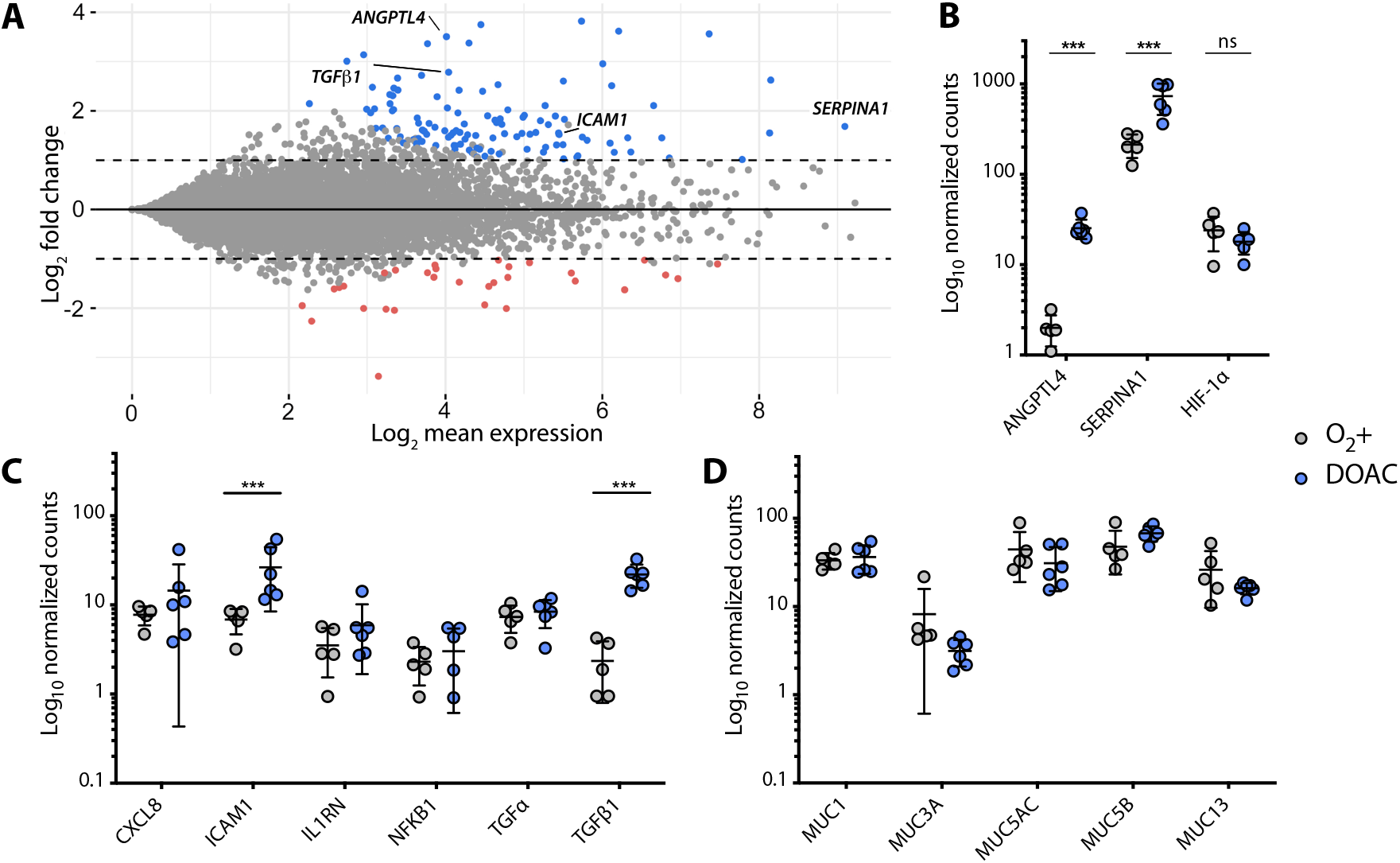
Dual oxic-anoxic culture of Calu-3 cells yields a similar transcriptomic profile to normoxic culture conditions (ALI). **(A)** MA plot representation of Calu-3 gene expression under DOAC conditions relative to normoxic culture at ALI (117 upregulated, 31 downregulated, log2fc > 1, padj <0.001). **(B)** *ANGPTL4* and *SERPINA1* were differentially expressed, though HIF-1α was consistent between cultures. Few changes in **(C)** inflammatory biomarkers and **(D)** mucin gene expression were observed. Data shown in panels B-D are log_10_-normalized gene counts from five or six independent biological replicates and were compared using the Wald test, Benjamini-Hochberg adjusted (***, *p*<0.001).

### Anaerobic airway microbiota promote an inflammatory host response

The lack of tractable epithelial cell culture systems compatible with hypoxic or anoxic bacterial growth has limited our understanding of host-anaerobe interactions. Prior work has shown that culture supernatants of anaerobic bacteria elicit pro-inflammatory cytokine expression *in vitro* through mixed-acid fermentation and production of SCFAs ^1,12,13^. However, it is not yet known how the host responds to the direct presence of anaerobes at the epithelial interface. To address this knowledge gap, we used the DOAC platform to assess the response of Calu-3 cells to co-culture with anaerobic microbiota.

As a starting point, we used a defined anaerobic bacterial consortium (ABC) derived from airway mucus collected from an individual with chronic sinusitis. This representative community was enriched under strict anaerobic culture conditions^15^ and was chosen for its dominant bacterial taxa (*Veillonella, Prevotella, Streptococcus*) associated with both healthy and diseased airways (**Fig. 3A**). These genera are also known for their mucin-degradation capacity and the ability to support pathogen growth through nutrient cross-feeding^14,15^. After 3h of equilibration under DOAC, Calu-3 cells were apically challenged with the anaerobically enriched bacterial consortium (∼8 × 10^6^ colony forming units (CFUs)) and incubated for an additional 24h. Colonization of the apical interface was confirmed and visualized using scanning electron microscopy (**Fig. 3B)**. Bacteria (3.6 × 10^6^ CFUs) were recovered after 24h by washing with PBS and plating on Brain Heart Infusion agar (BHI)(**Fig. 3C**), while washing with Triton X-100 resulted in a 0.8-log increase in recovery (2.4 × 10^7^ CFUs), suggesting both anaerobe growth at the epithelial surface and either robust attachment, disruption of bacterial aggregates, or bacterial invasion of host cells (i.e., cells were not removed by gentle PBS washing alone). Importantly, recovery of 8.1 × 10^5^ CFUs on *Prevotella* selective media under anoxic conditions (compared to 2.6 × 10^5^ in the inoculum) confirmed that apical oxygen concentrations were sufficiently low for strict anaerobe viability. We note that in contrast to the canonical airway pathogens *S. aureus* and *P. aeruginosa* ^26,27^, Calu-3 cytotoxicity was not induced by anaerobe treatment after 24h (**Fig. 3D**).

**Fig. 3.**
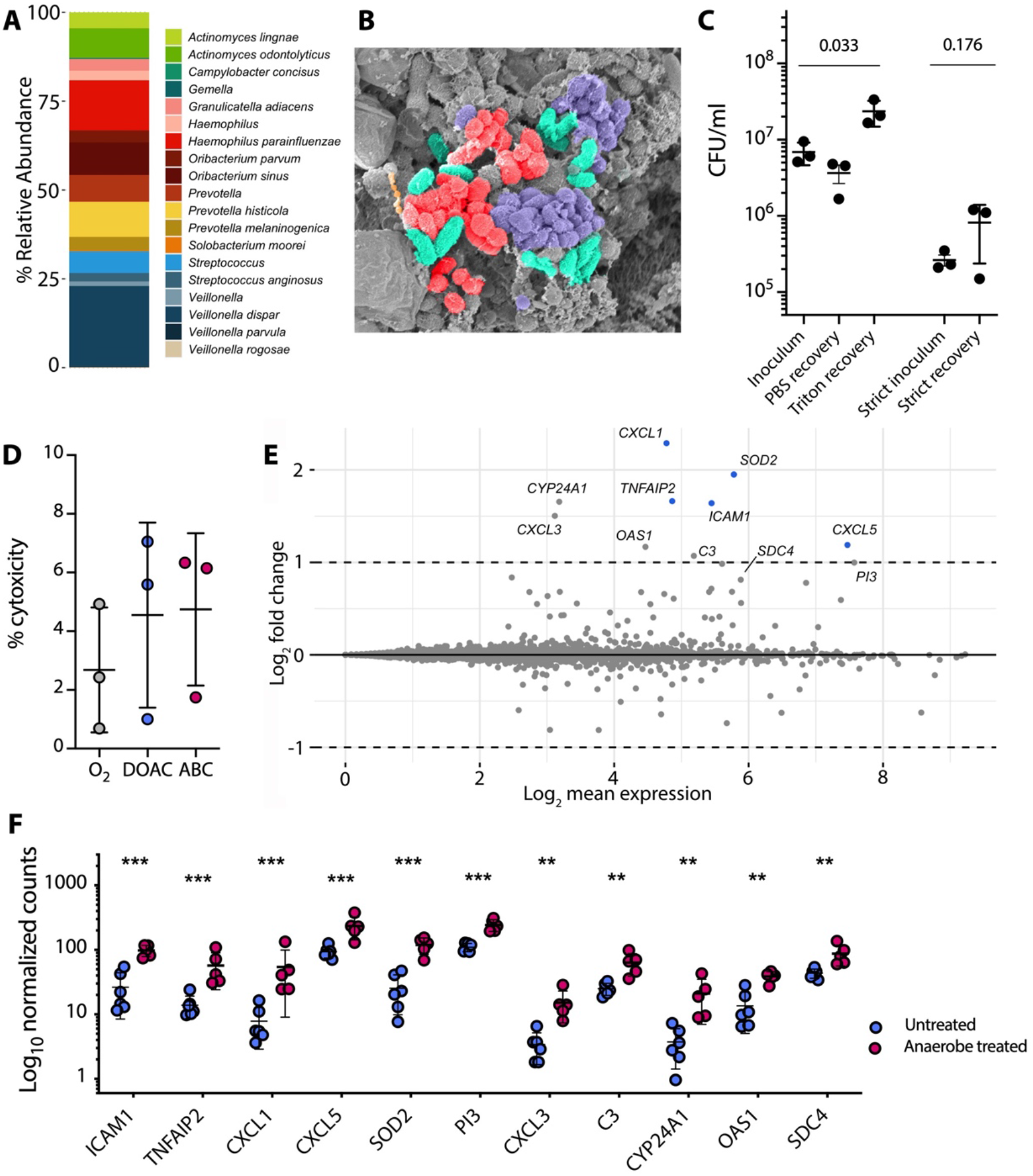
Anaerobic microbiota colonize the apical surface of Calu-3 cells and induce a pro-inflammatory response. **(A)** Taxonomic composition of an enriched anaerobic bacterial consortium (ABC) derived from the upper airways. **(B)** SEM micrograph of Calu-3 cells after challenge with ABC. **(C)** Bacterial recovery from Calu-3 cells after 24h by washing with PBS and TritonX-100. Strict anaerobe CFUs were determined by plate enumeration on Brucella Blood Agar under anoxic conditions. **(D)** Anaerobe (ABC) challenge did not result in Calu-3 cytotoxicity relative to unchallenged cells grown aerobically (O2) or under dual oxic-anoxic conditions (DOAC). **(E)** MA plot representation of Calu-3 gene expression under DOAC after ABC challenge relative to an untreated DOAC control. **(F)** Log_10_-normalized gene counts from five or six independent biological replicates. Data in panels C and D were compared using a one-way ANOVA with Dunn’s multiple comparisons test (n=3). Data in panels E and F were compared using a Wald test, Benjamini-Hochberg adjusted (\*\*\**p*<0.001, **<0.01).

We then used RNAseq to profile the Calu-3 transcriptional response to anaerobe challenge (**Fig. 3E, Data S2**) which revealed the differential expression of various inflammatory biomarkers. These included *ICAM1* (intercellular adhesion molecule 1, previously shown to be expressed in response to periodontopathic bacteria)^28^, *TNFAIP2* (mediated by TNFα in response to bacterial challenge)^29^, chemokines *CXCL1* and *CXCL5* (neutrophil chemoattractants primarily expressed as an acute inflammatory response to infection)^30,31^, and *SOD2* (superoxide dismutase 2, which is expressed in response to lipopolysaccharide and has an antiapoptotic role against inflammatory cytokines)^32^. Other inflammatory markers including *PI3* (elafin, an elastase inhibitor that can prime innate immune responses in the lung)^33^, *CXCL3* (neutrophil chemoattractant), *C3* (complement), *CYP24A1* (cytochrome p450 family 24 subfamily A member 1), *OAS1* (oligoadenylate synthetase), and *SDC4* (syndecan 4) were also differentially expressed, though did not reach our threshold adjusted *P* value of <0.001 (**Fig. 3F**). These data demonstrate that while not cytotoxic, anaerobes and/or their metabolic byproducts were detectable by host cells and elicit an inflammatory response.

### Anaerobic microbiota alter the mucosal interface via mucin degradation

Our previous work demonstrated the ability of anaerobic microbiota to degrade airway mucins and support the growth of canonical pathogens via nutrient cross-feeding^14,15^. Thus, in support of downstream pathogen colonization experiments, we used fast protein liquid chromatography (FPLC) to determine whether anaerobe challenge altered Calu-3 mucin integrity relative to unchallenged cells. To do so, we collected and purified mucin from the apical side of the Transwells as previously described^24^ and used size-exclusion chromatography to assay their integrity. As expected, chromatograms revealed two characteristic peaks; (i) high molecular weight mucins which ran in the void volume of the column, and (ii) a broader inclusion volume peak representative of lower molecular weight mucins^34^ (**Fig. 4A**). While differences in the chromatographic profile of peak 1 (high molecular weight mucins) were negligible between culture conditions (*P*=0.38), peak 2 area was significantly reduced (*P*=0.043) following anaerobe challenge (**Fig. 4B**), reflecting proteolytic and/or glycolytic degradation of lower-molecular weight mucin glycoproteins. These data suggest that in addition to eliciting a pro-inflammatory host response, anaerobic colonization can alter the physicochemical nature of the mucosal interface.

**Fig. 4.**
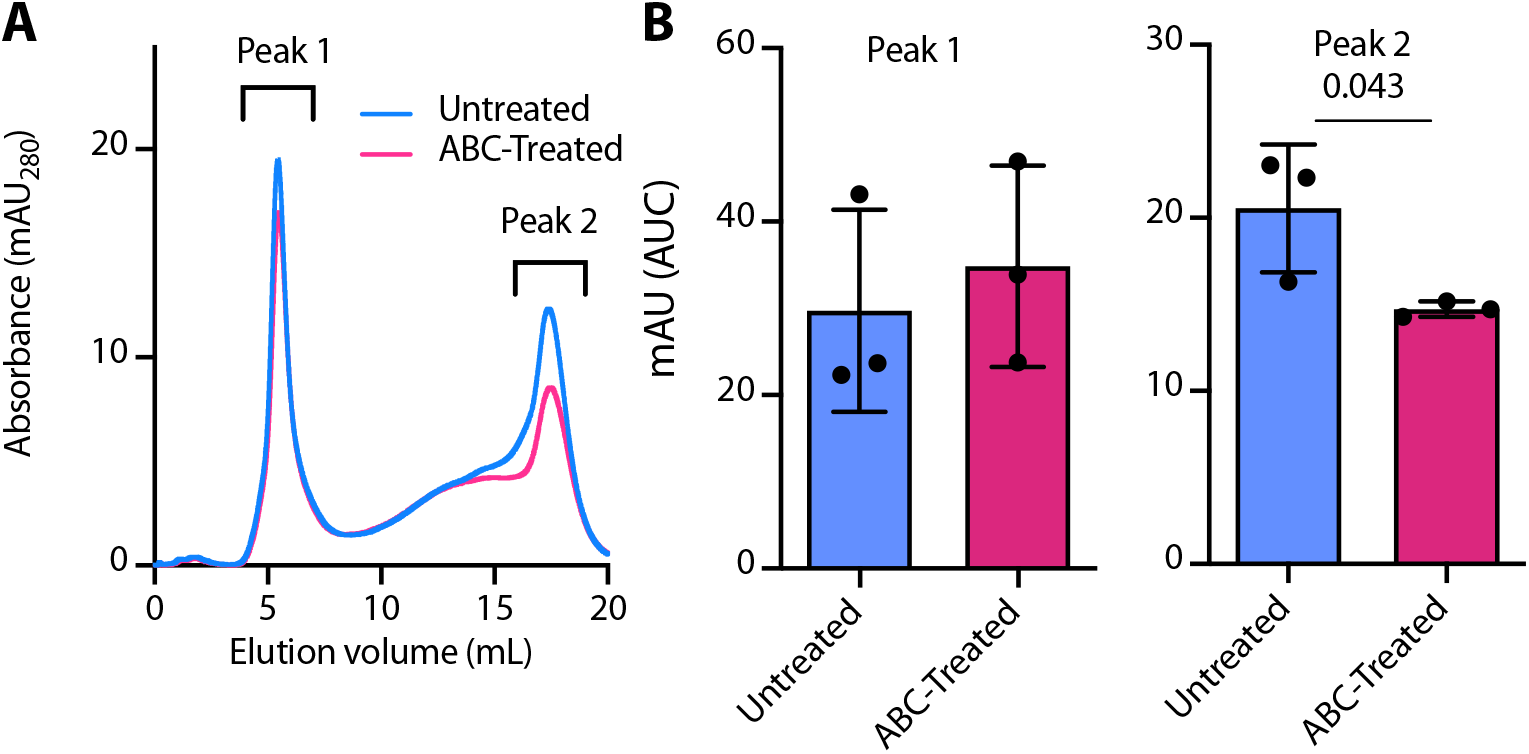
Anaerobic microbiota alter epithelial mucin integrity. **(A)** Representative FPLC traces of MUC5AC mucins purified from Calu-3 cells grown under DOAC conditions (untreated) and after treatment with an anaerobic bacterial community. **(B)** Area under curve (AUC) for both peak 1 (high molecular weight mucins) and peak 2 (low molecular weight mucin). Data shown were derived from three independent experiments using three biological replicates (n=9). Data were compared using a unpaired t-test with Welch’s correction (**, *p*<0.01).

### Anaerobes promote *P. aeruginosa* colonization of the airway epithelium

Viral challenge of the airway epithelium potentiates colonization by *P. aeruginosa* via interferon mediated effects^18^. Other work has shown that the protective role of the mucus barrier is compromised by *Streptococcus mitis* through hydrolysis of mucin glycans^35^. Given that the anaerobically enriched consortium used in our model both elicits inflammation (**Fig. 3**) and alters mucin integrity (**Fig. 4**), we hypothesized that in addition to providing nutrients for pathogen growth through cross-feeding, anaerobes enhance *P. aeruginosa* colonization of the airway epithelium (**Fig. 5A**).

**Fig. 5.**
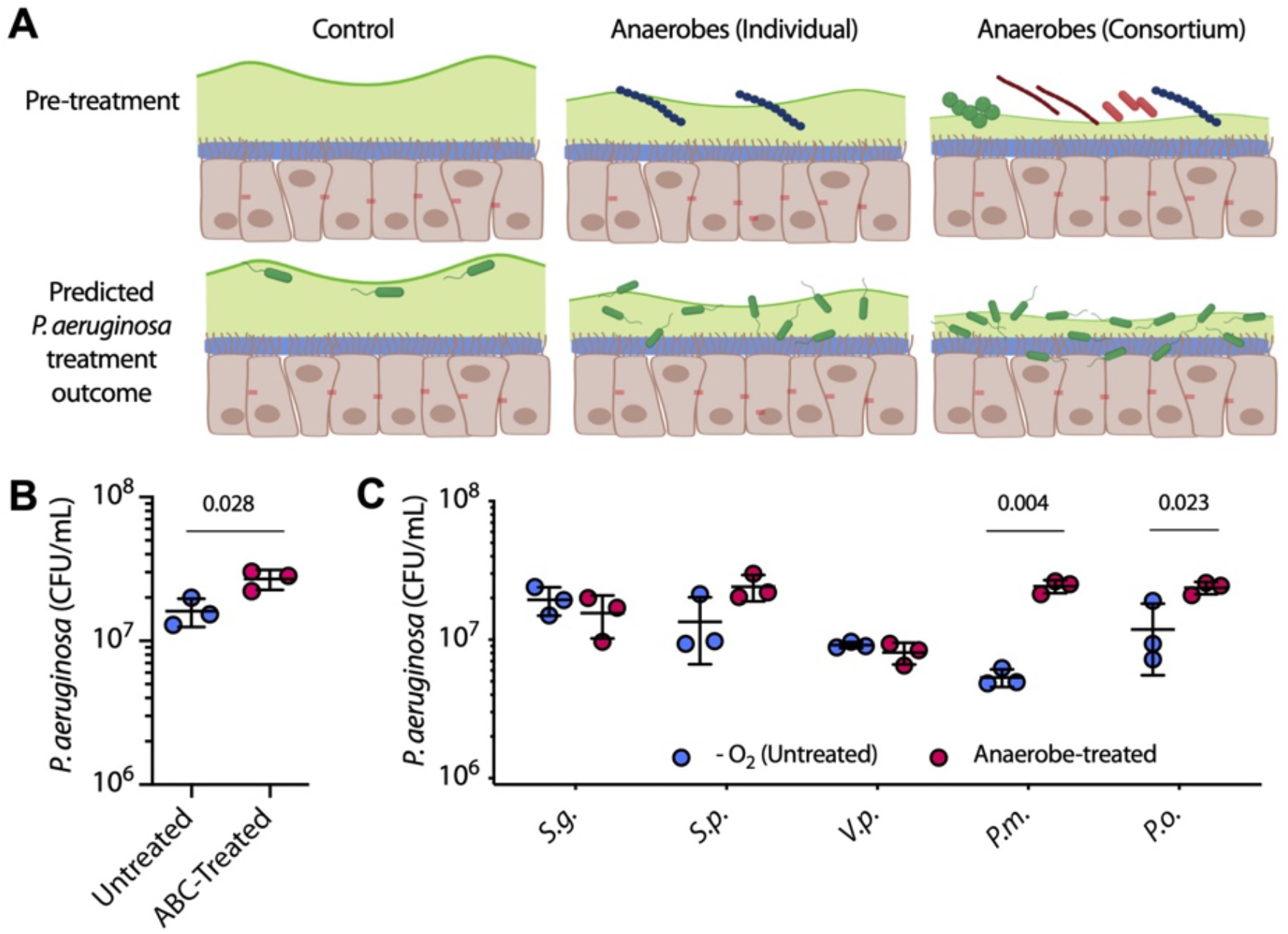
Anaerobic microbiota condition the epithelial interface for pathogen colonization. **(A**) Schematic of experimental design and expected outcomes. **(B)** Calu-3 pre-treatment with an anaerobic bacterial consortium (ABC-treated) potentiates *P. aeruginosa* colonization. **(C)** *P. aeruginosa* adhesion to Calu-3 cells after pre-treatment with individual anaerobes (*S*.*g*., *Streptococcus gordonii; S*.*p*., *S. parasanguinis; V*.*p*., *Veillonella parvula; P*.*o*., *Prevotella oris; P*.*m*., *P. melaninogenica)*. Data shown in panels B and C were derived from three independent experiments with three biological replicates (n=9). Pairwise comparisons in panel D were performed using an unpaired t-test with Welch’s correction.

To test this hypothesis, Calu-3 cells were first treated with the anaerobic consortium (see **Fig. 3A**) for 24h, washed with PBS to remove spent medium and unbound cells, and subsequently infected with ∼5 × 10^7^ CFUs of *P. aeruginosa* PA14 for 2h. Removal of spent medium and a short incubation time ensured that any difference in colonization between anaerobe-treated cells and an unconditioned (i.e., no anaerobe) control was a result of cell attachment and not enhanced growth over time. *P. aeruginosa* colonization was then determined by washing and permeabilization of Calu-3 cells followed by plate enumeration. As predicted, anaerobe pre-treatment resulted in an ∼67.5% increase in *P. aeruginosa* attachment (1.1 × 10^7^ CFUs) relative to untreated controls (*P*=0.028, **Fig. 5B**), supporting our hypothesis that anaerobic microbiota can enhance pathogen colonization of the airway mucosal interface.

To assess the contributions of individual anaerobes to *P. aeruginosa* colonization we then challenged Calu-3 cells with representative isolates of the three most abundant genera in the anaerobic consortium (*Streptococcus, Veillonella, Prevotella)* prior to *P. aeruginosa* colonization (**Fig. 5D**). We found that *Streptococcus* species (*S. gordonii* and *S. parasanguinis*) had little effect on PA14 colonization, despite their known mucolytic capacity. Similarly, *V. parvula*, an asaccharolytic bacterium, resulted in no significant differences between treatment conditions. By contrast, challenge with both *P. melaninogenica* and *P. oris*, two species commonly associated with inflammatory airway disease^1,6,36^, resulted in significantly increased PA14 recovery from Calu-3 cells compared to unconditioned controls (*P=*0.004 and *P=*0.023, respectively). These data suggest that while anaerobic microbiota of the respiratory tract facilitate enhanced colonization of the epithelium by *P. aeruginosa*, they do so in a species-specific manner.

### Single cell RNA sequencing reveals cell-specific expression of inflammatory marker genes and cancer-related pathways in response to *F. nucleatum*

A drawback to transcriptomic studies of epithelial cell populations (**e.g**., **Fig. 2A, Fig. 3E**) is that bulk RNA sequencing averages out contributions of different cell types to the overall transcriptional landscape. Not only does this mask how physiological processes are distributed among cells, but contributions from rare cell types are also largely invisible. To address these limitations, single-cell RNA sequencing (scRNAseq) has been widely adopted in respiratory research and has already led to the discovery of ionocytes, the immune landscape of lung cancer, novel mononuclear phagocytes in the CF lung, and the host response to SARS-CoV-2 infection^37-40^. To our knowledge, scRNAseq has not yet been applied to bacterial-host interactions in the context of airway disease. Given the many examples of tissue and cell-specific tropism exhibited by pathogens of the respiratory tract^41-43^, here we combined the DOAC platform with scRNAseq to test our hypothesis that anaerobic microbiota elicit an inflammatory host response that varies by epithelial cell type.

To test our hypothesis, primary human bronchial epithelial (nHTBE) cells were cultured under normoxia, transferred to DOAC conditions for 3h, followed by apical challenge with *Fusobacterium nucleatum* -both an anaerobic oral commensal and pathogen associated with a spectrum of respiratory tract infections^23^. After 24h of co-culture, we captured nHTBEs using the 10X Genomics Chromium controller **(Fig. 6A)** and sequenced an average of 2,007 cells per untreated replicate (n=6) and 3,682 from each *F. nucleatum-*treated replicate (n=3). After removing cells with less than 250 detectable genes and additional filtering criteria, we analyzed a total of 6,981 untreated and 5,010 treated cells, with an average of 4,042 and 3,103 expressed genes per cell, respectively. To identify main cell types, we performed a combined analysis of all cells from all samples using uniform manifold approximation and partitioning (UMAP)^44,45^. This analysis partitioned cells into 13 distinct clusters based on gene expression profiles, colored accordingly in **Fig. 6B**, which did not appreciably differ between treatments (**Fig. 6C; Fig. S3**). Clusters were manually identified post-hoc by identifying genes significantly upregulated within each cluster and by comparison to established sets of marker genes for canonical epithelial cell types (**Fig. 6D**)^37^. Among these clusters, we found cell type-specific expression of 17 pattern recognition receptors (PRRs) that play a critical role in the innate host response to microbial encounter (**Fig.6E; Fig.S3**). For example, toll-like receptor 2 (TLR2) was predominately expressed by secretory cells, while NOD1 and NLRP1 exhibited higher expression in ciliated and basal cells, respectively. These data are consistent with previous reports describing heterogeneous expression of PRRs across airway epithelial cell types^46,47^ and supported our hypothesis that the host response to bacterial challenge would also be cell type specific.

**Fig. 6.**
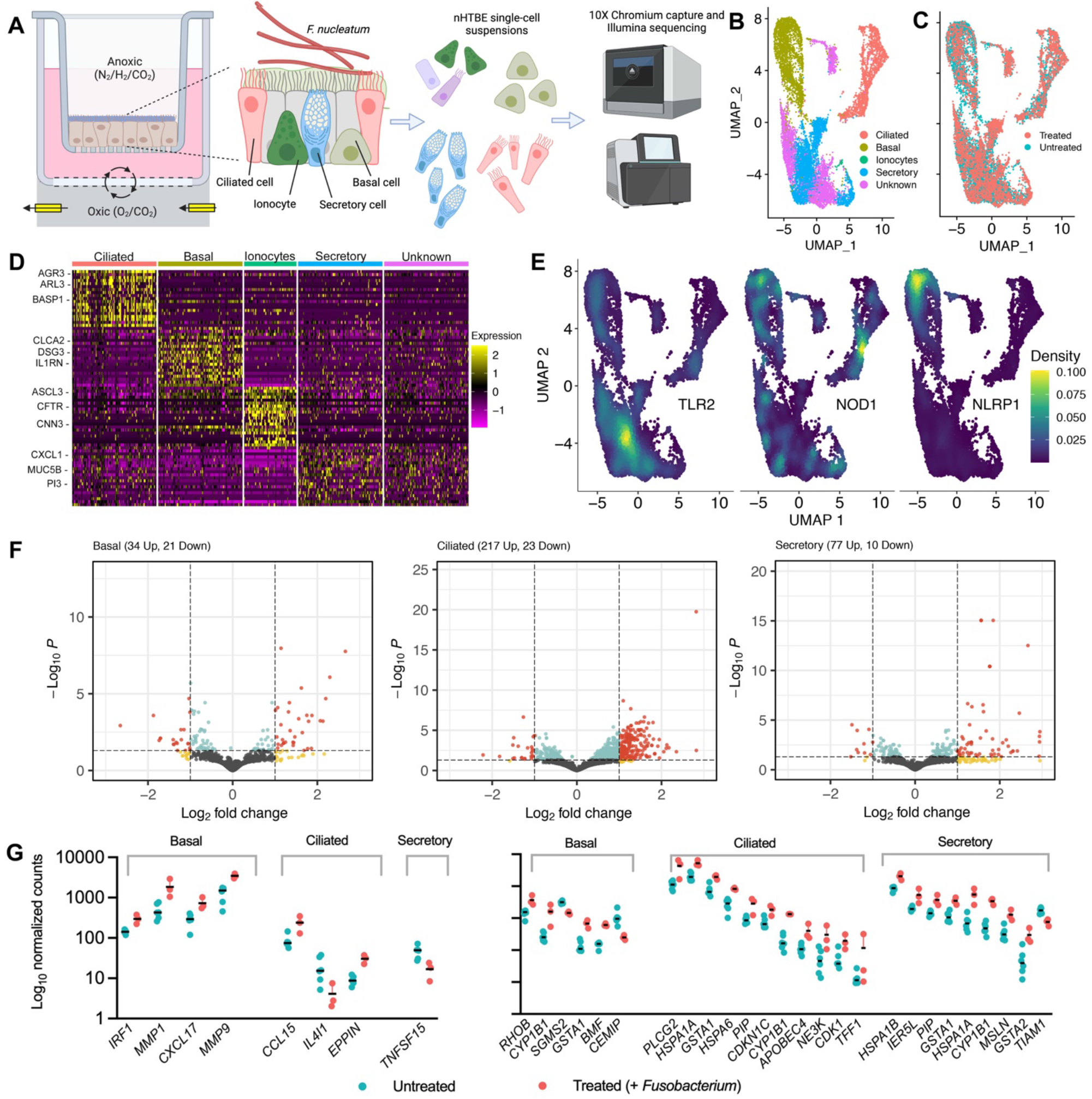
Single cell RNA sequencing. **(A**) Experimental schematic. Primary airway epithelial cells were cultured under DOAC conditions and challenged with *F. nucleatum*. Host cells were dissociated and collected via the 10X Genomics platform prior to scRNAseq. (**B**) UMAP projection of nHTBEs showing 4 major cell types (and unknown cells) from 13 distinct clusters (see Fig. S3). **(C)** UMAP projection of nHTBE cell types by treatment condition. **(D)** Scaled expression of the top DEGs that informed specific cell subsets. **(E)** Density plots (Nebulosa projections) of TLR2, NOD1, and NLRP1 showing heterogeneous expression of pattern recognition receptors between cell types. **(F)** Volcano plot representation of differential gene expression in basal, ciliated, and secretory cells between *F. nucleatum*-treated nHTBE cultures relative to untreated controls. **(G**,**H)** Scatterplot representations of differentially expressed (**G**) inflammatory markers (left panel) and cancer-associated genes (right panel) by cell type.

Using DESeq2^48^, we then performed differential gene expression analysis, accounting for both the effects of cell type and bacterial challenge. As predicted, comparison of *F. nucleatum-* treated cells to untreated controls revealed a distinct, cell type-specific response to bacterial challenge: basal, 34 up / 21 down; ciliated, 217 up / 23 down; and secretory cells, 77 up / 10 down (padj > 0.001) (**Fig. 6F, Data S3**). Among notable differentially expressed genes, *F. nucleatum* stimulated expression of multiple cytokines and chemokines, including tumor necrosis factor superfamily member 15 (*TNFSF15*) in secretory cells, *CCL15* in ciliated cells, and inflammatory marker genes encoding interferon regulatory factor (*IRF1*), matrix metalloproteases (*MMP1, MMP9*), and *CXCL17* in basal cells (**Fig.6G**). These data are consistent with previous reports of Fusobacteria stimulating inflammatory host responses in the oral cavity and diverse extra-oral body sites^49-52^. Unexpectedly, *F. nucleatum* challenge also resulted in a striking differential expression of cancer-associated genes and pathways (**Fig.6H)**. *CYP1B1*, a universal cancer marker encoding cytochrome P450 1B1^53^, and *GSTA1* (glutathione S-transferase A1) which promotes lung cancer cell invasion and adhesion^54^, both increased in expression in each of basal, ciliated, and secretory cells in response to *F. nucleatum* (**Fig. 6H)**. *HSPA1A* (encoding heat shock protein A), involved in the promotion of tumor cell proliferation, metastasis, and invasion^55^, was also upregulated in both basal and ciliated cells. Twenty-one other cancer-related genes varied by cell type (padj < 0.001). While *F. nucleatum* has been associated with late-stage lung cancer^56,57^ and poor patient response to immunotherapy^58^, these data support a direct role for the bacterium in airway carcinogenesis. To our knowledge, these data are the first to reveal the epithelial response to bacterial challenge with the granularity afforded by scRNAseq and further demonstrate the power and versatility of the DOAC platform to model anaerobe-host interactions.

## DISCUSSION

The surge in culture-independent sequencing of human microbiota has spawned renewed interest in the importance of anaerobic bacteria in disease etiologies. However, despite anaerobes comprising a significant component of airway bacterial communities, studies on their contributions to disease pathophysiology have reported seemingly contradictory results, calling into question their role(s) in patient morbidity. Proposed pathogenic mechanisms are supported by compelling *in vitro* data^12-14^, but their relevance is unresolved due to a lack of compatible models with which to test their interactions with the respiratory epithelium. To address this knowledge gap, here we optimized and characterized a model platform (DOAC) that enables co-culture of strict and facultative anaerobes with polarized airway epithelial cells. Importantly, provision of oxygen exclusively to the basolateral side of host cells prevented anoxia-associated cytotoxicity, inflammation, and significant changes in global gene expression, validating its utility for the study of anaerobe-host interactions. Using this platform, we demonstrate that (i) anaerobic microbiota promote enhanced epithelial colonization by *P. aeruginosa*, and (ii) that *F. nucleatum* elicits a pro-inflammatory and cancer-associated transcriptional response that varies by epithelial cell type.

Disparate oxygen demands between epithelial cells and anaerobic bacteria pose significant challenges for their co-culture *in vitro*. Several culture systems have been developed to overcome these challenges and recapitulate an oxygen-restricted mucosal interface^59-61^, though each has limitations. Transwell cultures of Caco-2, HaCaT, and primary human gingival cells have been used to demonstrate that anaerobic taxa can adhere to, invade, and alter oral and intestinal epithelia, yet these assays are either limited to short incubation times or poorly mimic *in vivo* conditions^62-64^. Newer microfluidic-based and ‘organ-on-a-chip’ models have also seen widespread interest due to their ability to establish a dual oxic-anoxic interface and facilitate study of anaerobe-host interactions^65-68^. However, these models either preclude direct host-bacterial contact or are technically challenging to maintain both oxic and strict anoxic microcompartments. The approach described here offers the distinct advantages of ease-of-use, direct interactions between host and microbiota, reproducibility given the multi-well plate format, and methodological flexibility lending itself to biochemical, microscopy, and transcriptomic studies.

To demonstrate the utility of the DOAC platform, we first used an anaerobic polymicrobial consortium representative of chronic airway disease. Dominated by *Streptococcus, Prevotella*, and *Veillonella* spp., this consortium (among other bacterial taxa) is thought to seed the airways through microaspiration from the oral cavity and is recognized as a key risk factor in the development of infection in COPD, CF, pneumonia, sinusitis and other diseases. For example, *in vitro* studies have shown that anaerobe-derived supernatants containing proteases and pro-inflammatory short-chain fatty acids modulate the immune tone of bronchial epithelial cell lines and primary cell cultures^12,13^. Indeed, anaerobe abundance in both diseased airways and healthy controls is associated with enhanced expression of inflammatory cytokines^1,9^. Our data also support an immunomodulatory role and suggest that anaerobe colonization of the airway mucosa may establish a local inflammatory environment known to promote colonization by *P. aeruginosa* and other canonical pathogens^18,20^.

We previously reported that anaerobes can also stimulate pathogen growth through mucin-based cross-feeding^14,15^. Specifically, *P. aeruginosa* and *S. aureus*, which cannot efficiently catabolize mucins in isolation, can gain access to bioavailable substrates via anaerobe-mediated degradation of the mucin polypeptide and O-linked glycans. Though not directly tested here, it is plausible that pre-colonization with an anaerobic bacterial community liberates additional mucin-derived metabolites on which pathogens can thrive at the epithelial interface. While it remains unclear why only low molecular weight mucins were altered (see Fig. 4), our FPLC data confirm that the mucosal surface is structurally modified because of anaerobic colonization. Given that mucin degradation has been shown to compromise its barrier function and enhance pathogen-epithelial interactions^35^, we propose that in addition to the host’s impaired mucociliary clearance, pathogenic contributions of anaerobic microbiota to airway infection are likely imparted through a multifactorial process (e.g., inflammation, cross-feeding, and surface alteration).

By focusing on an early time point after *P. aeruginosa* challenge (2h), we targeted anaerobic mucin degradation and its role in re-shaping the epithelial interface while enhancing pathogen attachment, as was previously shown for *S. mitis* and *Neisseria meningitidis*^35^. As predicted, anaerobe colonization resulted in a significant increase in *P. aeruginosa* attachment, likely mediated, at least in part, by mucin degradation. This observation has important clinical implications. Most notably, epithelial colonization is known to stimulate rapid *P. aeruginosa* biofilm maturation and associated increases in extracellular polysaccharide production, induction of quorum-sensing and other transcriptional changes^19^. In addition, *P. aeruginosa* grown on bronchial epithelial cells is far more resistant to antibiotic treatment than when grown on abiotic surfaces^19^, consistent with their increased tolerance *in vivo*. We propose that anaerobe-mediated colonization further potentiates these phenotypes. Moving forward, it will be important to consider *P. aeruginosa* interactions with the host over longer time periods to further understand differences in pathogen physiology and the inflammatory host response in the presence and absence of anaerobic microbiota.

We elected to test Calu-3 cells as a representative cell line for several reasons. First, Calu-3 cells reach polarization at ALI within ∼21 days and achieve TEER values far greater and more stable than those of primary cells (>1000 Ωcm^2^) ^69,70^. In addition, overproduction of mucus on the apical surface of Calu-3 cells mimics a diseased mucosal environment and allowed us to test the hypothesis that mucin degradation enhances pathogen colonization. This was an important consideration as *P. aeruginosa* biofilms, at least in the context of CF, are thought to form within secreted mucus as opposed to the epithelial layer ^71^. We also acknowledge the limitations of using Calu-3 cells. Unlike primary cells, which form a pseudostratified epithelium with mucociliary differentiation, Calu-3 cells are derived from human bronchial submucosal glands that comprise a relatively homogenous monolayer. Transcriptional and physiological responses to external stimuli may also be unique to Calu-3s. As an example, *S. aureus* enterotoxin B is known to elicit significant differences in barrier integrity as well as IL-6 and IL-8 production in Calu-3 cells relative to primary tissue^72^. These, among other considerations, underscore the importance of using the DOAC platform with additional cell lines and primary tissues to model anaerobe-host interactions in greater detail.

Indeed, the combined use of DOAC, primary bronchial epithelial cells, and scRNAseq revealed the cell-type specific transcriptional response to *F. nucleatum* challenge. Generally regarded as an abundant commensal of the oral cavity, there is growing recognition of *F. nucleatum* as an emerging pathogen associated with airway infection^15^, periodontal disease^73^, adverse pregnancy outcomes^74^, and GI disorders^75^. In the latter instance, colonic biopsies from patients with inflammatory bowel disease (IBD) carry a greater *F. nucleatum* burden, while strains isolated from inflamed IBD tissues tend to be more invasive than those from healthy controls^76,77^. Accumulating evidence also suggests its role in the progression of various cancers^78^, including those of the breast, oral cavity, esophagus, rectum, and colon, and data presented here likewise suggest a pro-inflammatory and tumorigenic effect of *F. nucleatum* on the airway epithelium. Recent work demonstrated that *Fusobacterium spp*., among other oral microbiota, correlate with lower airway inflammation and are enriched in advanced lung cancers^56,57^. However, to our knowledge, a direct effect of *F. nucleatum* on cancer-associated pathways in the airways was previously unknown, as was the cell type specific transcriptional response of the host. Whether *F. nucleatum* has a colonization preference for specific cells over others requires further study, but the unique distribution of PRRs between basal, secretory, and ciliated cells insinuates that detection of the bacterium and/or its secreted metabolites, is at least partially responsible for the heterogeneous host response. If immunomodulatory therapies are to be further developed and used clinically, understanding which bacterial species promote or mitigate inflammation / tumorigenesis and how individual cells respond will be critical information to have in-hand. The DOAC platform paves the way for further study of these and other aspects of the host-anaerobe dynamic.

In summary, DOAC represents a tractable co-culture system that facilitates extended interrogation of host-anaerobe interactions. While we use this model here to demonstrate a role for anaerobic microbiota in pathogen colonization of the airways and how the airway epithelium responds to oral-associated anaerobes, this work will undoubtedly benefit future studies focused on anaerobe-host and anaerobe-host-pathogen interactions. Not only do we anticipate generating a deeper understanding of our microbiota at the epithelial interface under oxygen-limited conditions, we also expect to identify new therapeutic strategies in addition to understanding how existing antimicrobials are impacted by hypoxic microenvironments known to exist *in vivo*. Finally, while we use an airway-derived bacterial community and *F. nucleatum* as our model organisms, this work motivates additional studies of the gut, lung, oral cavity, and other sites of infection, where etiological roles of anaerobes have been proposed but specific pathogenic mechanisms remain unclear.

## ACKNOWLEDGEMENTS

We thank the University of Minnesota Fabrication Shop for the custom design of our anaerobic chamber culture manifold. We are also grateful for Emma Stanley (10X Genomics) and Jerry Daniel at the University of Minnesota Genomics Center who provided assistance with scRNAseq. This work was supported by the National Heart Lung and Blood Institute (HL136919), Cystic Fibrosis Postdoctoral Fellowships to P.J.M. and J.R.F. (MOORE20F0, FLETC22F0), a National Institutes of Health fellowship (#T90DE0227232) awarded through the National Institute of Dental and Craniofacial Research to S.K.L., and an American Society for Microbiology Undergraduate Research Fellowship to L.A.K..

## AUTHOR CONTRIBUTIONS

P.J.M. and R.C.H. were responsible for study design and wrote the manuscript. P.J.M. T.D.W., L.A.K., S.J.A., S.K.L., J.R.F., and R.C.H., collected and analyzed the data and edited the manuscript. T.D.W., S.J.A., and S.K.L. processed and analyzed RNA sequencing data. S.M.O. contributed cell lines and assisted with experiments.

## STAR METHODS

### Epithelial Cell Culture

Calu-3 cells were maintained in Minimal Essential Medium (Corning, USA) in 10% fetal bovine serum (FBS, Gene) supplemented with 100 U/mL penicillin and 100 μg/mL streptomycin (Gibco) at 37°C in a 5% CO2 incubator. Upon reaching 80% confluency, 1 × 10^5^ cells were passaged onto 6.5mm culture inserts (24-well hanging inserts, 0.4 μm pore; Corning). When cells reached confluency (∼5 days), apical medium was removed to establish an air-liquid interface (ALI). Polarized cells were maintained for an additional 21-28 days, with basolateral media changes every other day, to facilitate differentiation and mucus accumulation.

Cell cultures were then assembled in a gas permeable multi-well plate manifold (**Fig. S2**). Briefly, once polarized, Calu-3 cells were placed into gaskets in a 24-well gas-permeable plate (CoyLabs, Grass Lake, MI) containing 800μl of MEM per well. Mineral oil (400μL) was added to unused wells to prevent gas permeation from the basolateral to apical side of the manifold. Once assembled and covered with a sterilized lid, the apparatus was moved to an anaerobic chamber (90% N2/5% H2/5% CO2) while mixed gas (21% O2/5% CO2/74% N2) was delivered to the base of the plate to oxygenate the basolateral side of the polarized monolayer. A schematic of this workflow is summarized in **Fig. S1**.

RPMI 2650 cells were similarly cultured, except Dulbecco’s Modified Eagle Medium (DMEM) was used as the basal growth medium. Primary nHTBE cells (Lonza Bioscience, CC-2540S) harvested from healthy individuals were expanded using Pneumacult-Ex Plus (Stem Cell Technologies) prior to seeding on 24-well inserts. Cells were maintained at ALI for ∼21 days in a Pneumacult ALI growth medium (Stem Cell Technologies) at 37°C and 5% CO2 in a humidified incubator.

### Cell culture assays

Calu-3 barrier integrity was determined by trans-epithelial electrical resistance (TEER) measured with a Millicell-ERS2 Volt-Ohm meter (Millipore Sigma). Barrier integrity was further assessed using the Human E-cadherin Quantikine enzyme-linked immunosorbent assay (ELISA) kit (R&D Systems, Minneapolis, MN). Cells were washed with 100μl of PBS, absorbance was measured at 450 nm using a BioTek Synergy H2 plate reader and concentrations of E-cadherin were determined against a standard curve according to manufacturer’s instructions. Tight junction formation was assayed using immunofluorescence with an anti-zonula occludens (ZO)-1 monoclonal antibody (Thermo). To do so, Calu-3 cultures on Transwell inserts were chemically fixed in 4% paraformaldehyde (PFA) in phosphate-buffered saline (PBS) for 1h at room temperature. Cells were blocked in 10% goat serum and 1% bovine serum albumin (Sigma) for 15 min. After blocking, cells were incubated with human anti-mouse ZO-1 (AlexaFluor-488 conjugate; ThermoFisher)(5 μg/mL) for 1h. Transwell membranes were washed, removed from the supporting plastic insert, and mounted on glass slides using Vectashield anti-fade mounting medium. Labeled cells were visualized (ex. 480nm, em. 525nm) on an Olympus IX83 inverted fluorescence microscope using a 20X objective lens (0.75 NA). Finally, cytotoxicity was determined by lactate dehydrogenase (LDH) measurements on cell-free supernatants collected from the apical compartment by washing with 100μL of PBS. LDH was quantified using the Cytotoxicity Detection Kit Plus assay (Roche) according to manufacturer’s instructions. Data were calculated as percent LDH release compared with a lysed control and reported as %LDH release = [(experimental value -low control)/(high control-low control)] x 100.

### Immunoassays

The inflammatory response of Calu-3 cells was measured after 24h of culture in the anaerobic chamber. To do so, spent medium was collected from the apical side of Transwell cultures, and interleukin (IL-) 8 and IL-6 were then measured by ELISA per manufacturer instructions (R&D systems). Untreated epithelial cells grown under standard incubator conditions (5% CO2) were used as a control.

### FPLC

Secreted mucins were collected from Calu-3 cells as previously described^24^. Briefly, cells grown on Transwell inserts were solubilized in a reduction buffer consisting of 6M guanidine hydrochloride, 0.1M Tris-HCl buffer, and 5mM EDTA (pH 8). Prior to solubilization, 10mM dithiothreitol (DTT) and a cOmplete Mini protease inhibitor tablet (Roche) were added to 400mL of reduction buffer to minimize mucin degradation. Cell suspensions were gently agitated by pipetting to dislodge biomass, and each of the six Transwell suspensions per plate were pooled into a single aliquot. Cells were rinsed with a reduction buffer to remove residual mucin. This mixture was then incubated for 5h at 37°C, followed by the addition of 25 mM iodoacetamide and incubation overnight at room temperature. Mucins were then dialyzed (1000 kDa MWCO) against 1L of 4M GuHCl buffer containing 2.25 mM NaH2PO4-H2O and 76.8 mM Na2HPO4 and proceeded for 36h with buffer exchanges every 12h.

Fast protein liquid chromatography (FPLC) size-exclusion chromatography was then used to evaluate the integrity of high-molecular weight mucins. Using an Akta Pure FPLC (GE Healthcare BioSciences, Marlborough, MA) housed at 4°C, 500 μL of purified mucin was manually injected and subjected to an isocratic run at a flow rate of 0.4 mL/min for 1.5 column volumes (CV) with 150mM NaCl in 50 mM phosphate buffer (pH 7.2) on a 15mL 10/200 Tricorn column packed with Sepharose 4B-CL beads. Data were collected using Unicorn 7 software (GE Healthcare Biosciences).

### Bacterial strains and culture conditions

*P. aeruginosa* PA14 was routinely cultured on Luria Bertani (LB) medium. *P. melaninogenica* ATCC 25845, *S. parasanguinis* ATCC15912, *V. parvula* ATCC10790, and *Fusobacterium nucleatum subsp. nucleatum* ATCC 25586 were obtained from Microbiologics (St. Cloud, MN). *S. gordonii* was obtained from M.C. Herzberg (University of Minnesota) and *P. oris* 12252T was purchased from the Japan Collection of Microorganisms. All anaerobes were maintained on Brain-Heart Infusion medium supplemented with hemin (0.25 g/L), vitamin K (0.025 g/L) and laked sheep’s blood (5% vol/vol) (BHI-HKB) in an anaerobic chamber. A mucin-enriched anaerobic bacterial community (ABC) derived from an individual with chronic rhinosinusitis was also used and was cultured in a minimal mucin medium (MMM)^14,15^. Community composition was determined using 16S rRNA gene sequencing as described previously ^15^.

### Bacterial challenge and infection

Forty-eight hours prior to bacterial challenge, Calu-3 cells were incubated in MEM + FBS without antibiotics. On the day of bacterial challenge, Transwells were assembled in the gas permeable culture system and transferred into the anaerobic chamber where they were equilibrated for 3h. Overnight cultures of each anaerobe and the anaerobic bacterial community (ABC) were grown in MMM. Each individual culture was diluted to a concentration of ∼1 × 10^6^ colony forming units (CFU) in MEM and 10 μL of bacterial suspension was added to the apical side of the Calu-3 cells. Similarly, 10 μL of an adjusted suspension (OD600nm = 0.1, ∼8 × 10^6^ CFUs) of the anaerobic community was added to separate wells. Co-cultures were then incubated for an additional 24h. Following anaerobe challenge, spent medium was collected and analyzed for cytotoxicity using the LDH colorimetric assay (described above). Mucins were also collected as described above for integrity analysis via FPLC. In a separate experiment, anaerobe viability was determined using plate enumeration. Briefly, Calu-3 cells were washed with 100μL of PBS (to remove loosely bound cells) or 0.25% Triton X-100 (to recover tightly bound or intracellular bacteria). Resulting washes were serially diluted and plated on BHI or Laked Brucella Blood Agar (BBA) with Kanamycin (100 μg/mL) and Vancomycin (7.5 μg/mL) for enumeration. The latter was used as the selective agents prevent growth of most obligate Gram-negative and Gram-positive anaerobic bacteria, including most facultative anaerobes.

For the *P. aeruginosa* colonization assay, Calu-3 cultures were removed from the anaerobic chamber following anaerobe challenge. Cells were gently washed with PBS and subsequently infected with 5 × 10^7^ CFU of *P. aeruginosa* for 2h. After incubation, cells were gently washed three times with PBS to remove unbound *P. aeruginosa* and were permeabilized using 0.25% Triton X-100. Bacteria were enumerated by plating serial dilutions of Calu-3 cell lysates on LB agar. All assays were performed using three biological replicates and data are reported as the mean of three experiments.

### Bulk RNA sequencing

The transcriptomic response of Calu-3 cells to anoxic culture and anaerobe challenge was determined using RNAseq. Calu-3 cells were cultured at ALI as described above and harvested after 24h of (i) normoxic growth without bacterial challenge, (ii) DOAC growth without bacterial challenge, and (iii) DOAC growth with anaerobic bacterial challenge. Normoxic, unchallenged cells were maintained under standard incubator conditions. At the conclusion of each experiment, RNAlater (Invitrogen) was added to the apical and basolateral side of each well. For each condition, RNA was isolated from at least 5 separate Transwells using the RNeasy Micro Plus kit (Qiagen) according to manufacturer’s instructions. DNase treatment was performed as part of the RNA Clean and Concentrator kit (Zymo). RNA quality (RIN > 9.7) and quantity were assessed using an Agilent Bioanalyzer and RiboGreen, respectively. cDNA libraries were prepared using the SMARTer Universal Low Input RNA Kit (Takara Bio) and submitted for sequencing at the University of Minnesota Genomics Center on the Illumina NovaSeq 6000 platform.

The Ensembl GTF annotation file was filtered to remove annotations for non-protein-coding features. Fastq files were evenly subsampled down to a maximum of 100,000 reads per sample. Data quality in fastq files was assessed with FastQC. Raw reads were mapped to reference Human (Homo_sapiens) genome assembly “GRCh38” using annotation from Ensembl release 98. Gene counts were generated with ‘featureCounts’ of the RSubread package^79^. DESeq2/1.28.1 was used to estimate size factors to generate normalized count data, estimate gene-wise dispersions, shrink estimates using type=‘ashr’, and perform Wald hypothesis testing ^48,80^. Genes with a log2 fold-change greater than 1 and Benjamini-Hochberg adjusted p-value < 0.001 were considered significant. Code and data files are shared at https://github.com/Hunter-Lab-UMN/Moore_PJ_2022 and NCBI Gene Expression Omnibus (GEO) Accession #GSE21824.

### Scanning electron microscopy

Co-cultures were washed three times in 0.2M sodium cacodylate buffer and submerged in primary fixative (0.15 M sodium cacodylate buffer, pH 7.4, 2% paraformaldehyde, 2% glutaraldehyde, 4% sucrose, 0.15% alcian blue 8GX) for 22h. Transwell membranes were washed three more times prior to a 90-minute treatment with secondary fixative (1% osmium tetroxide, 1.5% potassium ferrocyanide, 0.135M sodium cacodylate, pH 7.4). After three final washes, cells were dehydrated in a graded ethanol series (25%, 50%, 75%, 85%, 2 × 95%, and 2 × 100%) for 10 minutes each before CO2-based critical point drying. Transwell membranes were attached to SEM specimen mounts using carbon conductive adhesive tape and sputter coated with ∼5 nm iridium using the Leica ACE 600 magnetron-based system. Cells were imaged using a Hitachi S-4700 field emission SEM with an operating voltage of 2kV. Images were false colored using Adobe Photoshop CS6.

### Single cell RNA sequencing and analysis

nHTBE cells grown on Transwell inserts were incubated in antibiotic-free Pneumocult medium for 48h prior to bacterial challenge. nHTBEs were assembled in the DOAC system and transferred into the anaerobic chamber. After 3 hours of nHTBE equilibration, an overnight culture of *F. nucleatum subsp. nucleatum* ATCC 25586 grown in BHI was diluted to a concentration of 1 × 10^6^ CFU in Pneumacult medium and 10 μL of bacterial suspension was added to the apical side of the nHTBE cells in the anaerobic chamber. Co-cultures were incubated for 24 h prior to epithelial cell homogenization and cell capture.

Samples were prepared for scRNAseq as described previously^40^. Briefly, single cell suspensions were washed with PBS + 5% FBS, resuspended in 200μL of PBS + 5% FBS, passed through a 70 μm cell strainer, and placed on ice. Cells were counted using trypan blue on a BioRad TC20 automated cell counter. A targeted cell input of 5,000 cells per condition were used to generate Gel Bead-In Emulsion (GEMs). The Chromium Next GEM Single Cell 3’ Gel beads v3.1 kit (10X Genomics, Pleasanton, CA) was used to create GEMS following manufacturer’s instructions. Captured GEMS were used for cDNA synthesis and library preparation using the Chromium Single Cell 3’ Library Kit v3.1, followed by sequencing using the Illumina NovaSeq platform.

Raw count matrices were generated with Cell Ranger (v.3.0.1) for alignment to the human reference genome (Homo_sapiens.GRCh38). The resulting raw count matrix for each experimental data set was imported into an R pipeline using Seurat (v. 4.1.0)^81^ for quality control, normalization, integration, clustering, and differential expression analysis. Data were filtered to remove low quality cells using the following criteria: number of UMI counts per cell ≥ 500, number of genes per cell ≥ 250, number of genes detected per UMI ≥ 0.8, mitochondrial ratio less than 0.25, and genes expressed in ≥ 10 cells. ‘*SCTransform*’ was used to normalize the data and regress out variation due to mitochondrial gene expression. Cells were integrated across conditions using default parameters except all 300 most variable genes identified by *SCTransform* were used. Cell clustering was performed based on the first 20 principal components using the *FindClusters* function with a 0.4 resolution. Conserved markers in each cluster were identified using *FindConservedMarkers*. Cell types for each cluster were then identified using the top 10 markers by average log fold change across groups. This list of top markers was compared to known cell markers for basal cells, secretory cells, ciliated cells, and ionocytes (Plasschaert). Clusters with a minimum of two out of the ten top markers corresponding to a particular cell type were labelled as such. DESeq2 (v. 1.34.0) was then used to perform pseudobulk differential expression analysis across the different cell types accounting for *F. nucleatum* challenge. Specifically, this package was used to estimate size factors, normalized count data, estimate gene-wise dispersions, shrink estimates using type=‘apeglm’^82^, and perform Wald hypothesis testing. Genes with a log2 fold-change greater than 1 and Benjamini-Hochberg adjusted p-value < 0.05 were considered significant. Pattern recognition receptor gene expression was visualized in the untreated population using Nebulosa (v.1.4.0) *plot_density* function. All code and data files are shared at https://github.com/Hunter-Lab-UMN/Moore_PJ_2022.

