## Supplemental Figures 1-3 for "Dual oxic-anoxic co-culture enables direct study of host-anaerobe interactions at the airway epithelial interface"

**Moore et al.**

### Supplemental Data

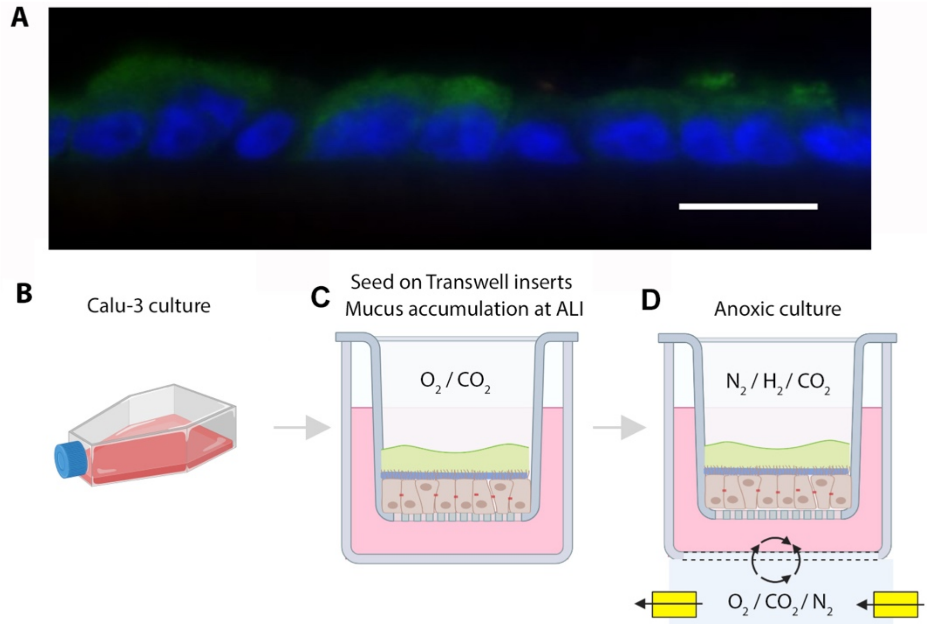


**Fig. S1. (A)** Confocal micrograph illustrating apical mucus production and accumulation by Calu-3 cells when cultured at air-liquid interface (anti-MUC5AC, green; DAPI, blue). Scale bar = 20 μm. **(B-D)** Dual oxic-anoxic culture (DOAC) of Calu-3 cells. **(B)** Calu-3 cells are cultured in MEM with 10% FBS. **(C)** Cells are seeded on 6.5mm Transwell inserts and grown to confluency (~5 days). Apical medium is removed, and cells are cultured at air-liquid interface (ALI) for 21-28 days under normoxic conditions prior to **(D)** incubation using the DOAC model where the apical compartment is oxygen limited and mixed gas is delivered basolaterally. Figure created with BioRender.com.

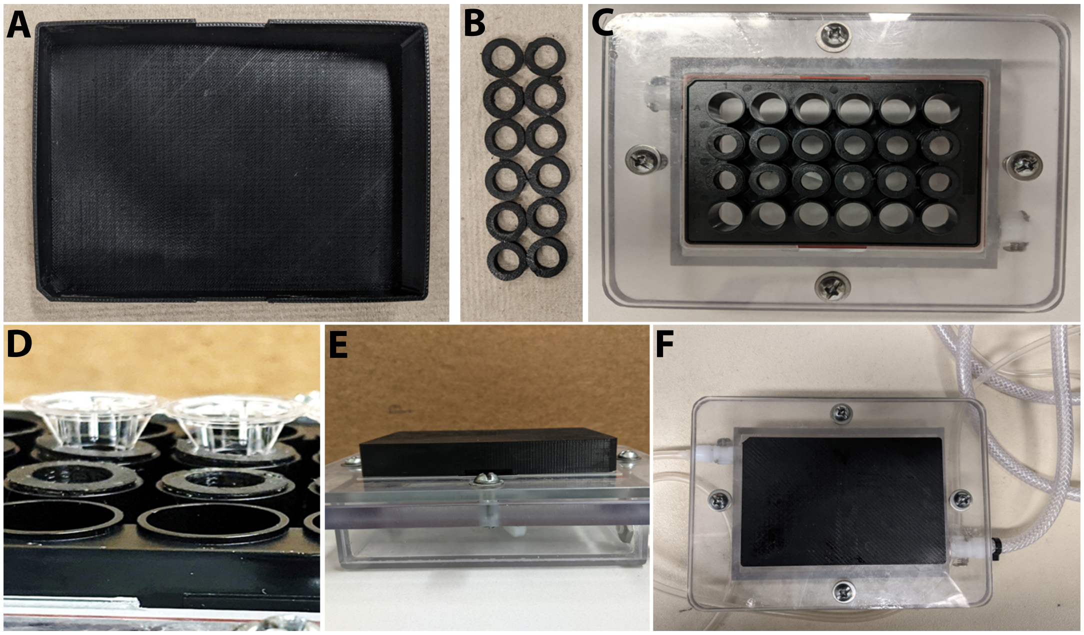


**Figure S2. Dual oxic-anoxic culture setup and methods.** Multi-well plate **(A)** lid and **(B)** gaskets were printed on a Stratasys F370 printer using thermoplastic polyurethane. SolidWorks files are attached as supplemental files (Supplemental File 1, File 2). A magnetic stir bar was placed in the base of console during assembly in a tissue culture hood. **(C)** A 24-well gas-permeable plate (Coy labs, Grass Lake, MI) was inserted into the plate base, followed by **(D)** sterile insertion of Transwell inserts containing polarized Calu-3 cells. 800 μL of DMEM was added to each experimental well, while 400 μL of sterile mineral oil was added to empty wells to prevent gas permeation. **(E)** Plates were covered and transferred to a Coy anaerobic chamber (90% $N_2$/5% $H_2$/5% $CO_2$). **(F)** Mixed gas (21% $O_2$/5% $CO_2$/74% $N_2$) was delivered and removed through inlet/outlet ports in the base plate. The system was allowed to equilibrate for 3h prior to bacterial challenge.

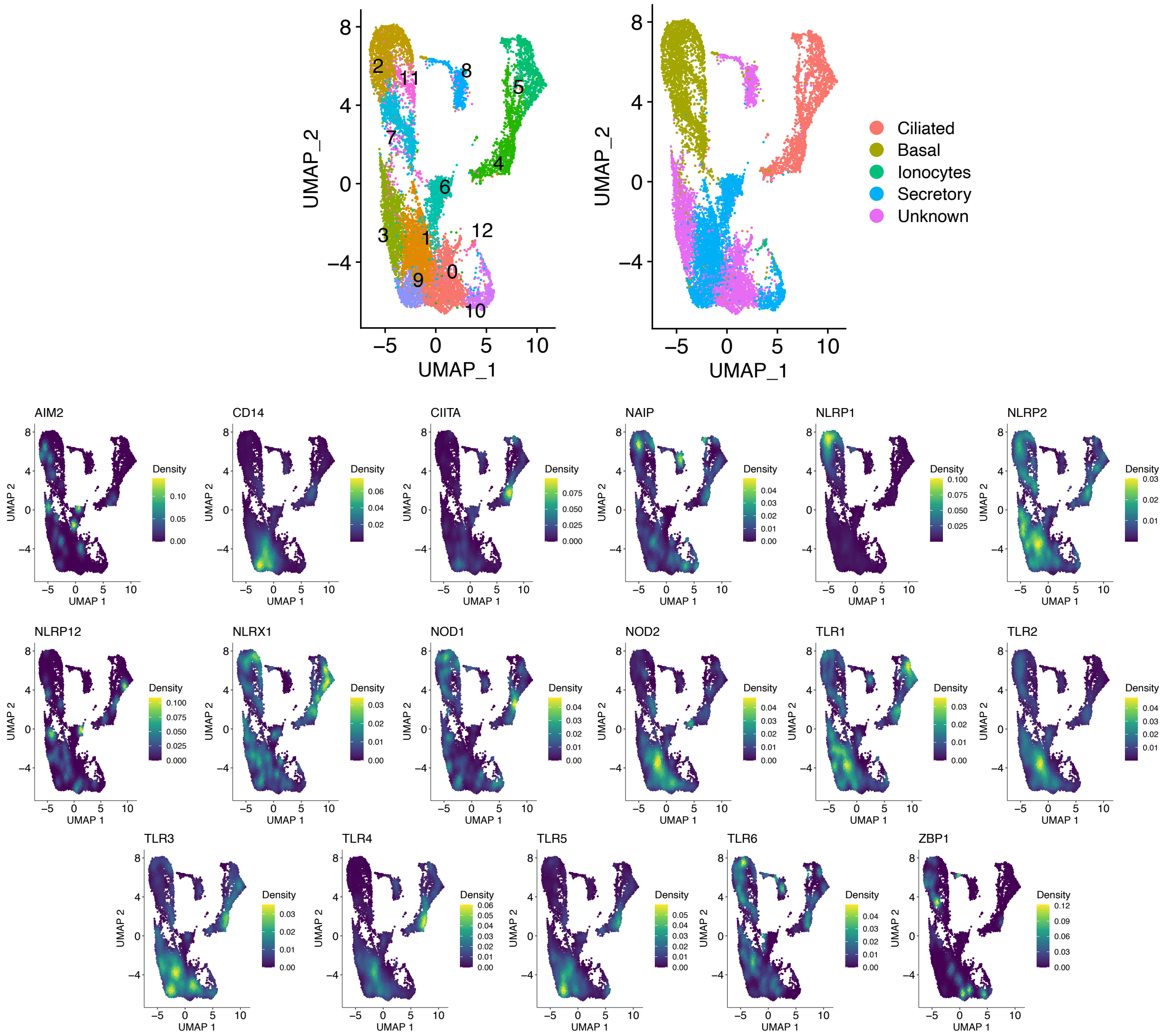


**Figure S3. Heterogeneous expression of pattern recognition receptors by epithelial cell type. (A)** UMAP projection of nHTBEs showing 13 distinct clusters based on global gene expression patterns. **(B)** Cell identity of each cell cluster was determined based on known marker genes. **(C)** Nebulosa plots reveal the cell-type specific expression patterns of Toll-like receptors (TLRs), nucleotide-binding oligomerization domain (NOD) receptors, NOD-Leucine Rich Repeat-containing receptors (NLR), and other pattern recognition receptors (PRRs) on nHTBEs.
